# GluN2D-containing NMDA receptors support dentate granule cell excitability, synaptic plasticity, and memory

**DOI:** 10.64898/2026.03.06.710109

**Authors:** Coralie Berthoux, Alma Rodenas-Ruano, Laxman Bist, Kaoutsar Nasrallah, Maryann Castillo, Gajanan P Shelkar, Shashank M Dravid, Sharon A Swanger, Pablo E Castillo

**Affiliations:** Dominick P. Purpura Department of Neuroscience, Albert Einstein College of Medicine, Bronx, NY 10461, U.S.A; Department of Biological Sciences, Fordham University, Bronx, NY 10458, U.S.A.; Department of Psychiatry and Behavioral Science, Texas A&M University, College Station, TX 77845, U.S.A.; Fralin Biomedical Research Institute at Virginia Tech Carilion, Roanoke, VA, U.S.A.; Department of Psychiatry & Behavioral Sciences, Albert Einstein College of Medicine, Bronx, NY 10461, U.S.A

**Keywords:** LTP, excitatory transmission, GluD1, hippocampus, learning, memory

## Abstract

N-Methyl-D-aspartate ionotropic glutamate receptors (NMDARs) are crucial for synaptic transmission, long-term plasticity, neuronal activity, and cognition. Consistent with these functions, NMDAR dysfunction is linked to several brain disorders, including Alzheimer’s disease, autism, schizophrenia, and depression. NMDARs are tetrameric complexes composed of two essential GluN1 subunits and two distinct GluN2 subunits (GluN2A-D) that define their functional characteristics. Although the roles of GluN2A and GluN2B, which are highly expressed in the brain, have been extensively studied, much less is known about GluN2D in brain function. Using selective GluN2D antagonists in the mature rodent brain and a conditional GluN2D knockout model, we assessed the role of GluN2D-containing NMDARs in dentate granule cells (GCs). We found that these receptors are tonically active, primarily extrasynaptic, and facilitate GC action-potential firing. Additionally, physiologically relevant presynaptic and postsynaptic activity patterns induced strong long-term potentiation of NMDAR-mediated transmission at medial perforant path synaptic inputs. This plasticity was likely driven by lateral diffusion of GluN2D and supported by non-canonical glutamate delta-1 (GluD1) receptors. Finally, removing GluN2D from excitatory cells in the dentate gyrus impaired spatial memory. Overall, our findings demonstrate that GluN2D-containing NMDARs are vital for hippocampal function, likely by modulating GC activity and mediating NMDAR synaptic plasticity.

**TEASER:** GluN2D-containing NMDA receptors control dentate gyrus function by regulating neuronal firing and synaptic plasticity

## INTRODUCTION

NMDA receptors (NMDARs) are crucial for brain development and function, playing key roles in synaptic plasticity, learning, and memory. Their dysfunction is linked to conditions such as Alzheimer’s disease, schizophrenia, depression, autism, and neurodevelopmental disorders (*1, 2*). NMDARs are tetrameric complexes typically composed of two GluN1 and two GluN2 subunits (GluN2A, GluN2B, GluN2C, and GluN2D). The GluN2 subunit subtypes confer distinct physiological and pharmacological properties to the receptor complex. In the adult forebrain, the GluN1/GluN2A/GluN2B triheteromer is the most abundant NMDAR subtype (*3*), and for this reason, GluN2A and GluN2B subunits have been widely characterized. Although the GluN2D subunit is also expressed in the mature brain (*4–6*), its role is largely understudied compared with other subunits. Understanding the specific contribution of GluN2D-containing NMDA receptors is important, as GluN2D subunits may contribute to excitotoxicity (*7, 8*), and mutations and polymorphisms associated with schizophrenia and infantile epilepsy have been identified in the *GRIN2D* gene, which encodes GluN2D (*9, 10*).

GluN2D mRNA expression in rodents is high during brain development, declines postnatally, and persists in certain brain areas, including the hippocampus, cerebellum, and basal ganglia (*5, 11–13*). Specific cell types in these areas express GluN2D-containing NMDARs that mediate synaptic transmission, and the tonic activity of these receptors increases neuronal firing (*14–23*). Functional studies indicate that GluN2D is expressed extrasynaptically in most neurons, including CA1 pyramidal cells (*21*) and interneurons (*16, 24, 25*). In situ hybridization in human tissue has revealed *GluN2D* mRNA in dentate granule cells (GCs) (*26*), and a study in rats using GluN2D-preferring antagonists suggests the presence of functional GluN2D-containing receptors in these neurons (*27*). However, the role of these receptors in hippocampal function remains unknown.

The dentate gyrus plays a crucial role in learning and memory by transforming patterns of information from the entorhinal cortex into distinct neuronal representations and transferring them to area CA3 (*28–31*). GCs are the most abundant excitatory neurons in the dentate gyrus and convey the only output from this region via mossy fibers, which are the axons of GCs. GCs receive excitatory inputs from the medial and lateral entorhinal cortex (MEC and LEC) via the medial and lateral perforant paths (MPP and LPP), respectively (*32*). Activity-dependent plasticity and dendritic integration are essential for information processing in the dentate gyrus (*33–36*). MPP inputs onto GCs exhibit robust AMPAR long-term potentiation (LTP) (*37–39*). Furthermore, early studies showed that high-frequency stimulation can also trigger NMDAR plasticity at MPP-GC excitatory synapses (*40, 41*). Whether GluN2D-containing receptors contribute to GC function and activity-dependent plasticity is unclear.

Here, we investigated the role of the GluN2D subunit in GCs of the mature rat and mouse brains using selective GluN2D antagonists and a complementary genetic approach to conditionally delete *Grin2d* from GCs. We report that GluN2D-containing receptors regulate action potential firing in GCs, mediate long-term synaptic plasticity, and are essential for spatial memory.

## RESULTS

### Dentate granule cells express functional extrasynaptic GluN2D-containing NMDA receptors

GluN2D expression in the rodent brain declines postnatally, especially in glutamatergic neurons (*5, 11–13*). To address whether GCs express *Grin2d* mRNA in the mature mouse brain, we used single-molecule fluorescence in situ hybridization (FISH) in P18 and P45 mice (**Figure 1A**). Parvalbumin interneurons, which express high levels of *Grin2d* mRNA in the hippocampus (*42*), served as a positive control. We found that *Grin2d* was expressed in the granule cell layer of the mature mouse brain*. Grin1,* which encodes the NMDAR obligatory subunit GluN1, was also highly expressed in GCs, as expected. Most cells in the granule cell layer that expressed *Grin2d* did not express *parvalbumin* mRNA. Together, our results indicate that *Grin2d* is expressed in GCs of the mature mouse dentate gyrus, strongly suggesting the presence of GluN2D-containing NMDARs in these neurons.

**Figure 1.**
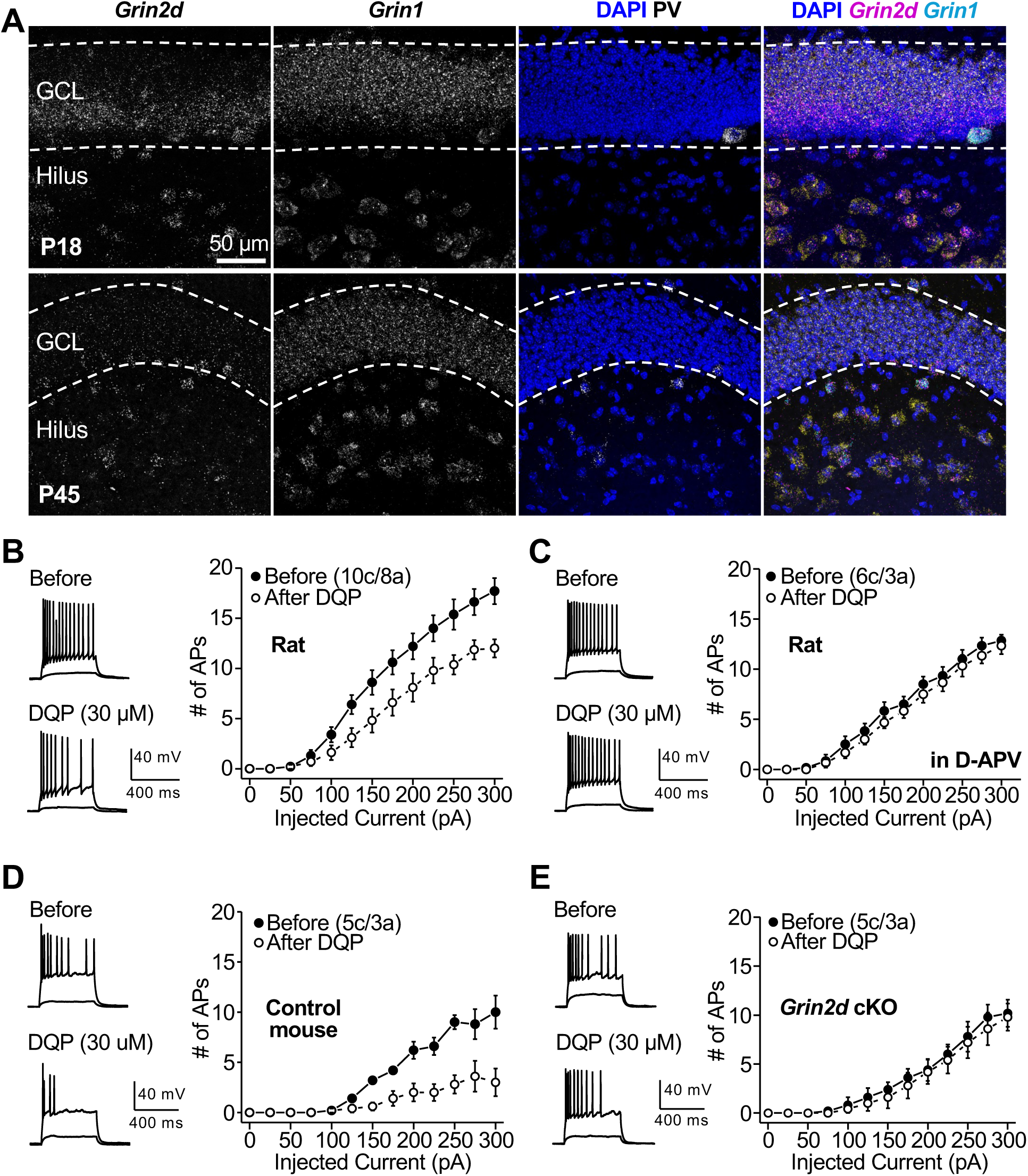
GluN2D-containing NMDARs tonically control GC firing. **(A)** *Grin2d* mRNA is expressed in dentate granule cells (GCs). Representative 63x images show *Grin1*, *Grin2d*, and *parvalbumin* RNA in the ventral dentate gyrus of P18 and P45 mice. Nuclei are labeled with DAPI. *Grin2d* is expressed throughout the granule cell layer (GCL), with the most prominent expression in cells near the subgranular zone. *Grin1* mRNA is expressed in all neurons in the GCL and hilus. *Parvalbumin* labeling shows that most cells expressing *Grin2d* in the GCL and subgranular zone are not interneurons, indicating that *Grin2d* is expressed in GCs. **(B)** Representative traces (*left*) and summary plot (*right*) showing that bath application of the GluN2D antagonist DQP-1105 (30 μM for 15 min) significantly decreased GC firing elicited by direct current injection (p < 0.001, n = 10, two-way ANOVA with Tukey’s comparison test) in rat hippocampal slices. **(C)** DQP-induced decrease in firing was abolished in the presence of NMDAR antagonist D-APV (50 μM) (p = 0.2154, n = 7, two-way ANOVA with Tukey’s comparison test). **(D)** Representative traces (*left*) and summary plot (*right*) showing that GluN2D antagonist DQP-1105 (30 μM) caused a significant decrease in GC firing in control mice (p < 0.001, n = 5, two-way ANOVA with Tukey’s comparison test). **(E)** This effect was absent in *Grin2d* cKO mice (p = 0.22976, n = 5, two-way ANOVA with Tukey’s comparison test). Data are presented as mean ± s.e.m. Numbers in parentheses refer to the number of recorded cells (c) and animals (a) for this and all remaining figures. Representative traces correspond to the firing induced by a 250-pA current injection and the voltage change elicited by a subthreshold current (∼ 25 pA) in the presence of 10 μM NBQX and 100 μM picrotoxin.

Tonic NMDAR activation by ambient glutamate (for a recent review, see *43*) can facilitate neuronal firing in CA1 pyramidal neurons (*44, 45*), dentate GCs (*46*), prefrontal cortical pyramidal cells and fast-spiking interneurons (*47*), interneurons in CA1 stratum radiatum (*48*), and chick auditory neurons (*49*). This action requires the GluN2D subunit, at least in inhibitory interneurons (*18, 24, 50, 51*). To determine whether GluN2D-containing NMDARs tonically regulate GC activity, we tested the effect of the non-competitive GluN2C/GluN2D-selective antagonist DQP-1105 (*52, 53*) on GC firing. Although this antagonist may not distinguish between GluN2C- and GluN2D-containing NMDARs, GluN2C expression in the adult hippocampus (*54*) and in neurons in general (*55*) is negligible. We found that bath application of 30 μM DQP-1105 for 15 min reduced action potential firing in GCs evoked by current injection in rat hippocampal slices in the presence of 10 µM NBQX and 100 µM picrotoxin to block AMPAR- and GABA_A_ receptor-mediated synaptic transmission, respectively (**Figure 1B**). This effect was absent when DQP-1105 was applied in the presence of the NMDAR antagonist D-APV (50 μM) (**Figure 1C**), indicating that NMDARs are required for the DQP-1105-mediated reduction of GC firing. We then tested whether this reduction was mediated by GC GluN2D-containing NMDARs by examining *Grin2d* conditional knock-out (cKO) mice. We injected an AAV_5_-CaMKII-Cre-mCherry into the dentate gyrus of floxed *Grin2d* mice (*Grin2d^fl/fl^*) mice, while animals injected with AAV_5_-CaMKII-mCherry served as controls (**Figure S1**). DQP-1105 also reduced GC excitability in mouse hippocampal slices (**Figure 1D**), and this effect was abolished in *Grin2d* cKO mice (**Figure 1E**). Moreover, bath application of DQP-1105 (30 μM for 15 min) significantly altered the holding current in GCs (holding potential [Vh] = +40 mV) (**Figure 2A,B**), suggesting a tonic activity mediated by GluN2D-containing NMDARs. In support of this possibility, the DQP-1105-mediated effect was abolished in the presence of 50 μM D-APV in the rat (**Figure 2A**) and in *Grin2d* cKO mice (**Figure 2B**). Altogether, these results strongly suggest that GCs in both rats and mice express tonically active GluN2D-containing NMDARs that facilitate action potential firing, consistent with previous findings in other brain areas as indicated above.

**Figure 2.**
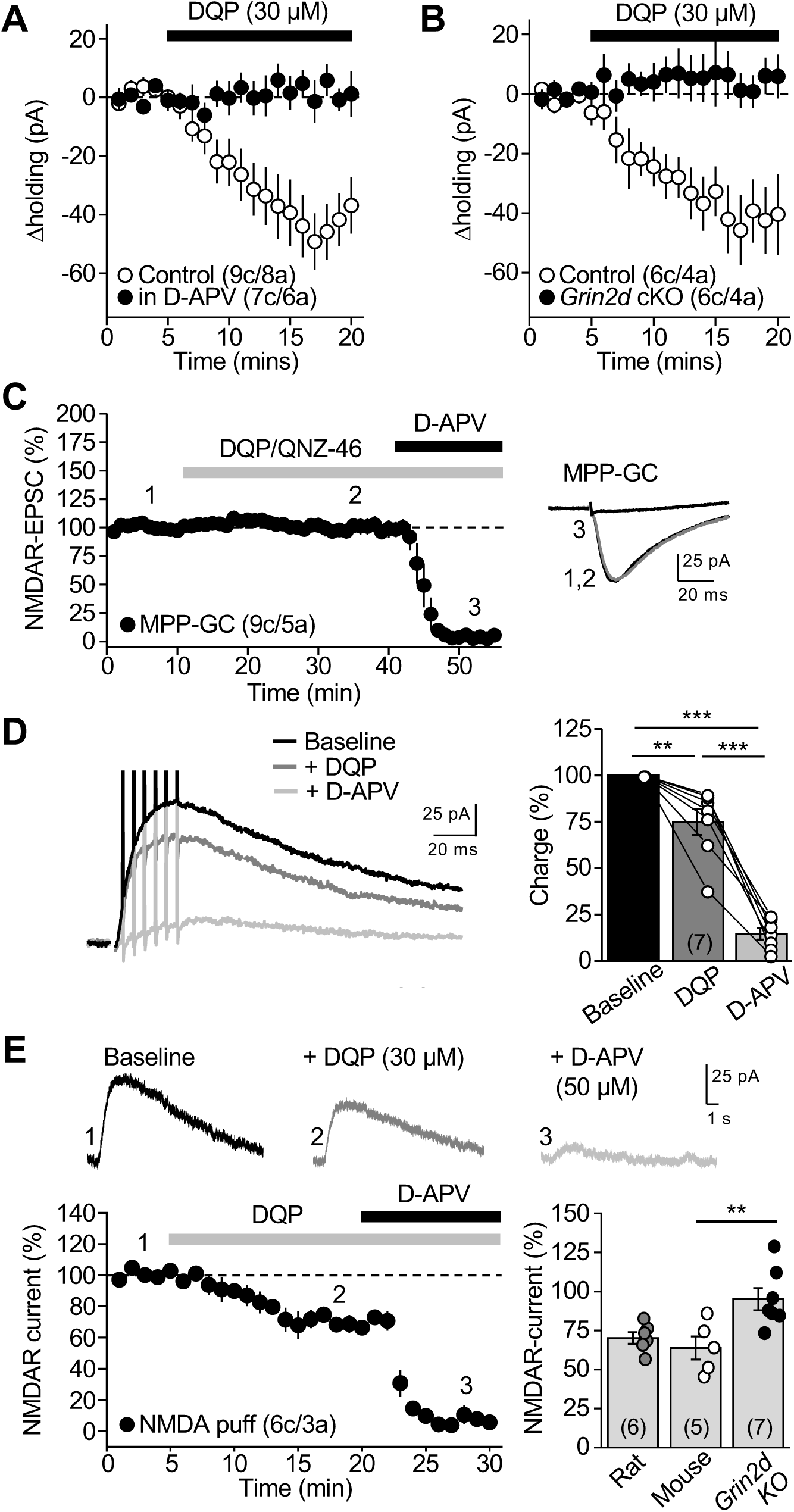
Pharmacological evidence for extrasynaptic GluN2D-containing NMDARs. **(A)** Bath application of the GluN2D antagonist DQP-1105 (30 μM for 15 min) caused a significant decrease in GC holding current (Control: −43.5 ± 1.3 pA, p < 0.01, n = 9, paired t-test), which was abolished in the presence of NMDAR antagonist D-APV (50 μM) (D-APV: 4.9 ± 4.7 pA, p = 0.3371, n = 7, paired t-test; Control *vs* D-APV: p < 0.001, unpaired t-test). Voltage-clamp recordings of GCs were performed at Vh = +40 mV in the presence of 10 μM NBQX and 100 μM picrotoxin. **(B)** The DQP-induced decrease in holding current was abolished *Grin2d* cKO mice (cKO: −0.4 ± 5.2 pA, p = 0.99479, n = 6, paired t-test) compared with controls (Control: −42.0 ± 11.5 pA, p < 0.05, n = 6, paired t-test; Control *vs* cKO: p < 0.01, unpaired t-test). **(C)** MPP NMDAR-EPSCs were recorded from GC (Vh = −45 mV) and evoked by stimulation electrode placed in the middle molecular layer (MML), in the presence of 10 μM NBQX and 100 μM picrotoxin. Bath application of the GluN2D antagonists DQP-1105 (30 μM) or QNZ46 (30 µM) for 30 mins had no effect on basal NMDAR synaptic transmission at MPP-GC synapses (99.9 ± 5.9 %, p = 0.9705, n = 9, paired t-test). The presence of NMDARs at the synapse was confirmed by bath applying D-APV (50 μM for 15 min) at the end of the recording. **(D)** *Left*, superimposed representative NMDAR-EPSCs elicited by repetitive stimulation of MPP inputs (7 stimuli at 200 Hz) before and after bath application of DQP-1105 (30 μM for 15 min) and subsequent application of D-APV (50 μM). *Right*, summary plot showing the effects of DQP and D-APV on the charge of the burst-induced NMDAR-EPSC (DQP-1105: 74.9 ± 7.1 %, p < 0.01 *vs* baseline; D-APV: 14.6 ± 3.1 %, p < 0.001 *vs* baseline, n = 7, one-way ANOVA with Tukey’s comparison test). **(E)** *Top*, representative NMDA outward currents (Vh = +40 mV) elicited by puffing a mix of 1 mM NMDA and 100 μM glycine on the MML of rat hippocampal slices. Bath application of the GluN2D antagonist DQP-1105 (30 μM) reduced the amplitude of these currents, which were abolished by subsequent application of D-APV (50 µM). *Bottom left*, time-course plot (DQP-1105: 70.2 ± 3.7 % of baseline, p < 0.001, paired t-test; D-APV: 6.4 ± 2.3 % of baseline, p < 0.001, paired t-test; n = 6). *Bottom right,* summary plot showing the effects of DQP in rats, wild-type (WT: 63.9 ± 7.4 %, p < 0.01, n = 5, paired t-test), and *Grin2d* KO mice (95.1 ± 7.1 %, p = 0.5022, n = 7, paired t-test; Control *vs* KO: p < 0.01, unpaired t test). Data are presented as mean ± s.e.m.

Although most evidence indicates that GluN2D-containing NMDARs are primarily extrasynaptic in adult animals, these receptors could also contribute to synaptic transmission (*1*). We therefore examined this possibility in GCs. To isolate NMDAR-mediated excitatory postsynaptic currents (NMDAR-EPSCs), whole-cell patch-clamp recordings of GCs (Vh = −45 mV) were performed in the presence of 10 µM NBQX and 100 µM picrotoxin. NMDAR-EPSCs were evoked by electrical stimulation with a patch-type pipette placed in the medial molecular layer (MML) of the rat dentate gyrus (see Experimental Procedures). Bath application of GluN2C/GluN2D-selective antagonists DQP-1105 (30 µM) or QNZ46 (30 µM) had no effect on NMDAR-EPSCs elicited by a single stimulation of MPP inputs onto GCs, whereas subsequent application of the non-selective NMDAR antagonist D-APV (50 µM) abolished MPP-evoked EPSCs (**Figure 2C**). We next tested whether glutamate spillover induced by repetitive stimulation of MPP inputs (7 stimuli at 200 Hz) could activate extrasynaptic GluN2D-containing NMDARs in GCs (*21*). Bath application of DQP-1105 (30 µM for 15 min) significantly reduced NMDAR-EPSCs (**Figure 2D**), suggesting that repetitive presynaptic activity may activate extrasynaptic GluN2D-NMDARs in GCs. In addition, we locally puffed NMDA (1 mM) and glycine (100 µM) onto the MML while voltage-clamping these cells at +40 mV. NMDA-elicited currents were significantly reduced by DQP-1105 (30 µM), and subsequent bath application of D-APV (50 µM) abolished these currents (**Figure 2E**). Lastly, puffing NMDA onto the MML of mouse hippocampal slices induced NMDA-mediated currents in GCs, which were significantly reduced by bath-applied DQP-1105, and this reduction was abolished in *Grin2d* KO mice (**Figure 2E**). Thus, although GluN2D-containing NMDARs in GCs do not significantly contribute to single-pulse-evoked basal MPP-GC synaptic transmission, they can be activated by short bursts of presynaptic activity and by exogenous NMDA application, strongly suggesting that GCs express functional extrasynaptic GluN2D-containing NMDARs.

### GluN2D is essential for burst timing-dependent NMDAR-LTP at MPP-GC synapses

We next examined whether GluN2D-containing receptors could contribute to long-term NMDAR plasticity at MPP-GC synapses in response to burst activity patterns. Burst timing-dependent plasticity (BTDP) of NMDAR-mediated transmission has been reported at other synapses (*56–58*). Therefore, we paired presynaptic (MPP axons) and postsynaptic (GC) burst activity to mimic *in vivo* activity in the entorhinal cortex (*59, 60*) and the dentate gyrus (*61, 62*). After monitoring a 10-minute stable NMDAR-EPSC baseline, we delivered a presynaptic (6 pre-pulses at 50 Hz)–postsynaptic (5 post-pulses at 100 Hz) burst-firing pairing protocol with a 10-ms interval, repeated 100 times, in current-clamp mode (Vh = −60 mV) (**Figure 3A,B**). We found that pre-post burst stimulation induced robust NMDAR-LTP at MPP-GC synapses, whereas a post-pre pairing sequence had no long-term effect on NMDAR-mediated synaptic transmission (**Figure 3C**). In addition, presynaptic and postsynaptic bursts alone did not elicit any NMDAR plasticity (**Figure 3D**).

**Figure 3.**
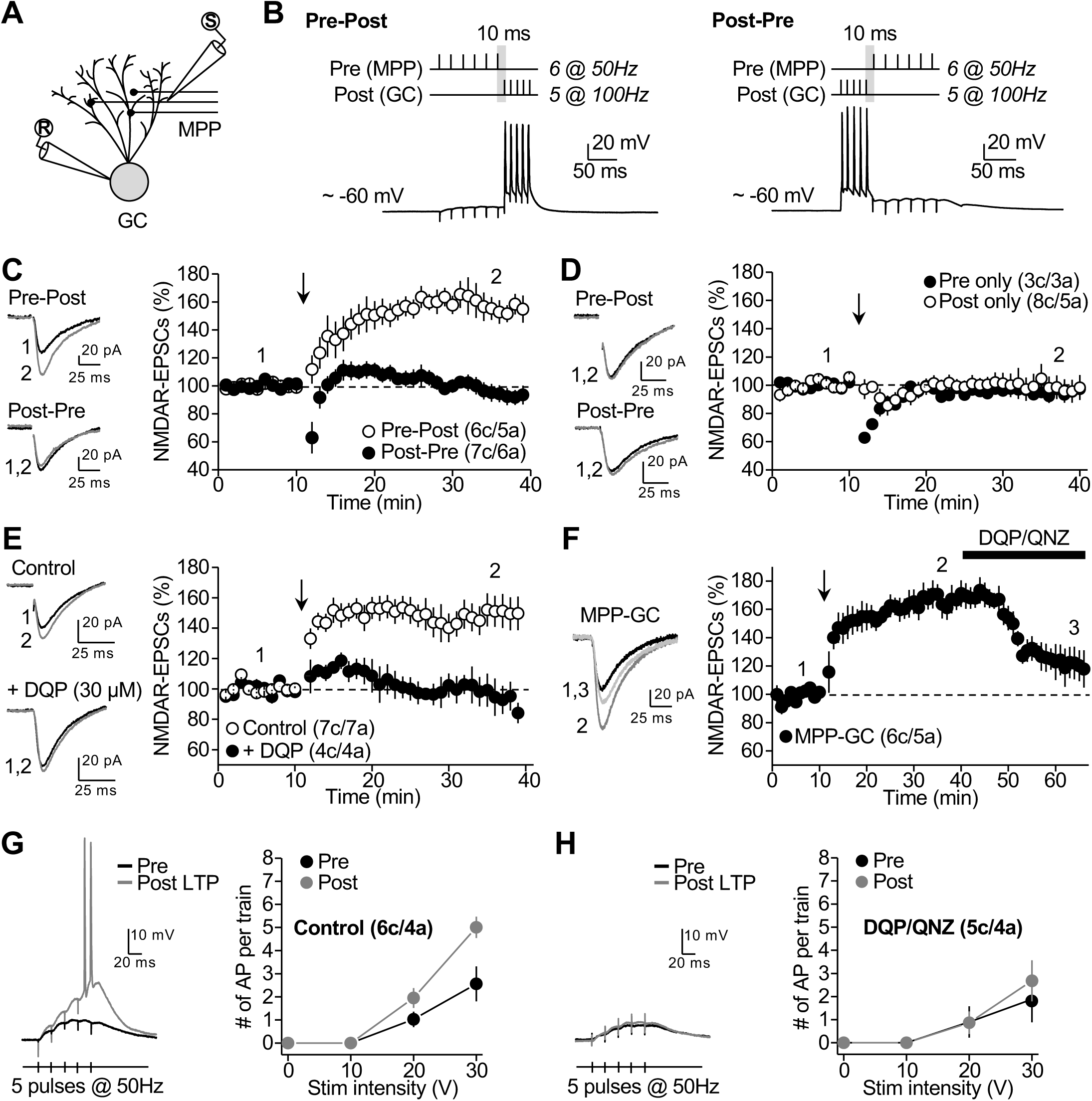
MPP-GC synapses express burst timing-dependent NMDAR-LTP. **(A)** Schematic diagram illustrating the recording configuration. Whole-cell patch-clamp recordings were performed from granule cells (GCs), and the stimulation electrode was placed in the middle molecular layer to stimulate medial perforant path (MPP) inputs. **(B)** Schematics of the induction protocols and example traces of single pairing. The pre-post protocol consisted of 6 pre pulses at 50 Hz followed by 5 post pulses at 100 Hz with a 10-ms interval, repeated 100 times at 0.5 s intervals. The post-pre protocol consisted of 5 post pulses at 100 Hz, followed by 6 pre pulses at 50 Hz, with a 10-ms interval, repeated 100 times at 0.5 s intervals. **(C)** *Left*, representative average traces before (1) and after (2) pairing protocol application. *Right*, time-course summary plot showing how pairing pre- and postsynaptic activity induced robust NMDAR-LTP at MPP-GC synapses in rat hippocampal slices (Pre-Post: 159.5 ± 7.3 %, p < 0.001, n = 6, paired t-test). In contrast, Post-Pre protocol did not induce significant long-term changes in NMDAR-mediated transmission (Post-Pre: 96.9 ± 5.3 %, p = 0.58, n = 7, paired t-test). The pairing protocol was delivered at the time indicated by the vertical arrow. NMDAR-EPSCs were recorded (Vh = −45 mV) in the presence of 10 μM NBQX and 100 μM picrotoxin. **(D)** Neither presynaptic (Pre only: p = 0.51, n = 3, paired t-test) nor postsynaptic (Post only: p = 0.89, n = 8, paired t-test) bursts alone elicited any long-lasting change in NMDAR EPSC amplitude. **(E)** Bath application of the GluN2D antagonist DQP-1105 (30 µM) prevented NMDAR-LTP induction (Control: 149.1 ± 8.1 %, p < 0.001, n = 7, paired t-test; DQP-1105: 95.6 ± 8.8 %, p = 0.65, n = 4, paired t-test; DQP-1105 *vs* control: p < 0.01, unpaired t-test). **(F)** Pre-post pairing LTP induction protocol was delivered at MPP inputs. Bath application of either DQP-1105 (30 µM) or QNZ46 (30 µM) 30 min after NMDAR-LTP induction substantially (166.9 ± 8.2 %) significantly reduced NMDAR-mediated transmission (119.5 ± 8.8 %, n = 6; GluN2D antagonism *vs* post-LTP: p < 0.05, unpaired t-test). **(G)** *Left,* example traces of whole-cell current-clamp recordings in a GC (Vh –60 mV) in response to MPP burst-stimulation (5 pulses at 50 Hz, 20 V), before and 15–20 min after induction of NMDAR LTP in the continuous presence of 10 µM NBQX, 100 µM picrotoxin, and 20 µM SCH50911 to block AMPA/kainate, GABA_A_, and GABA_B_ receptors, respectively. *Right*, average number of action potentials (APs) per train at different stimulation intensities, showing that induction of NMDAR LTP increases MPP-driven GC firing under control conditions (p < 0.001, n = 6, two-way ANOVA with Tukey’s comparison test). **(H)** This effect was abolished in the presence of DQP-1105 (30 µM) or QNZ46 (30 µM) (p = 0.42, n = 5, two-way ANOVA with Tukey’s comparison test). Data are presented as mean ± s.e.m.

To assess the role of GluN2D subunits in NMDAR-LTP, we first delivered the BTDP protocol in the presence of DQP-1105 (30 µM), which prevented NMDAR-LTP (**Figure 3E**). Moreover, bath application of DQP-1105 or QNZ46 (30 µM) 30 minutes post-induction –i.e., once LTP was established– significantly reduced LTP magnitude (∼73%), suggesting that most strengthening of NMDAR-EPSCs is mediated by GluN2D-containing receptors (**Figure 3F**). While NMDAR-EPSCs mediated by diheteromeric GluN2D-containing receptors exhibit a characteristic slow decay, the EPSC decay was not changed after LTP (**Figure S2**), suggesting that the recruited NMDA receptors could be triheteromeric complexes (e.g., GluN1/GluN2A/GluN2D or GluN1/GluN2B/GluN2D) (*3*).

NMDAR-LTP can enhance synaptically driven firing at other synapses (*63*). Therefore, we examined whether NMDAR-mediated transmission could enhance synaptically driven GC firing following LTP induction in a GluN2D-dependent manner. Brief activation of MPP inputs (5 stimuli, 50 Hz) significantly increased the number of action potentials per burst (**Figure 3G**), and this effect was abolished in the presence of DQP-1105 or QNZ46 (30 µM) (**Figure 3H**). Lastly, NMDAR-LTP was also induced in control mouse hippocampal slices but was abolished in *Grin2d* cKO mice (**Figure 4A**), indicating that postsynaptic GluN2D-containing NMDARs are necessary for NMDAR-LTP.

**Figure 4.**
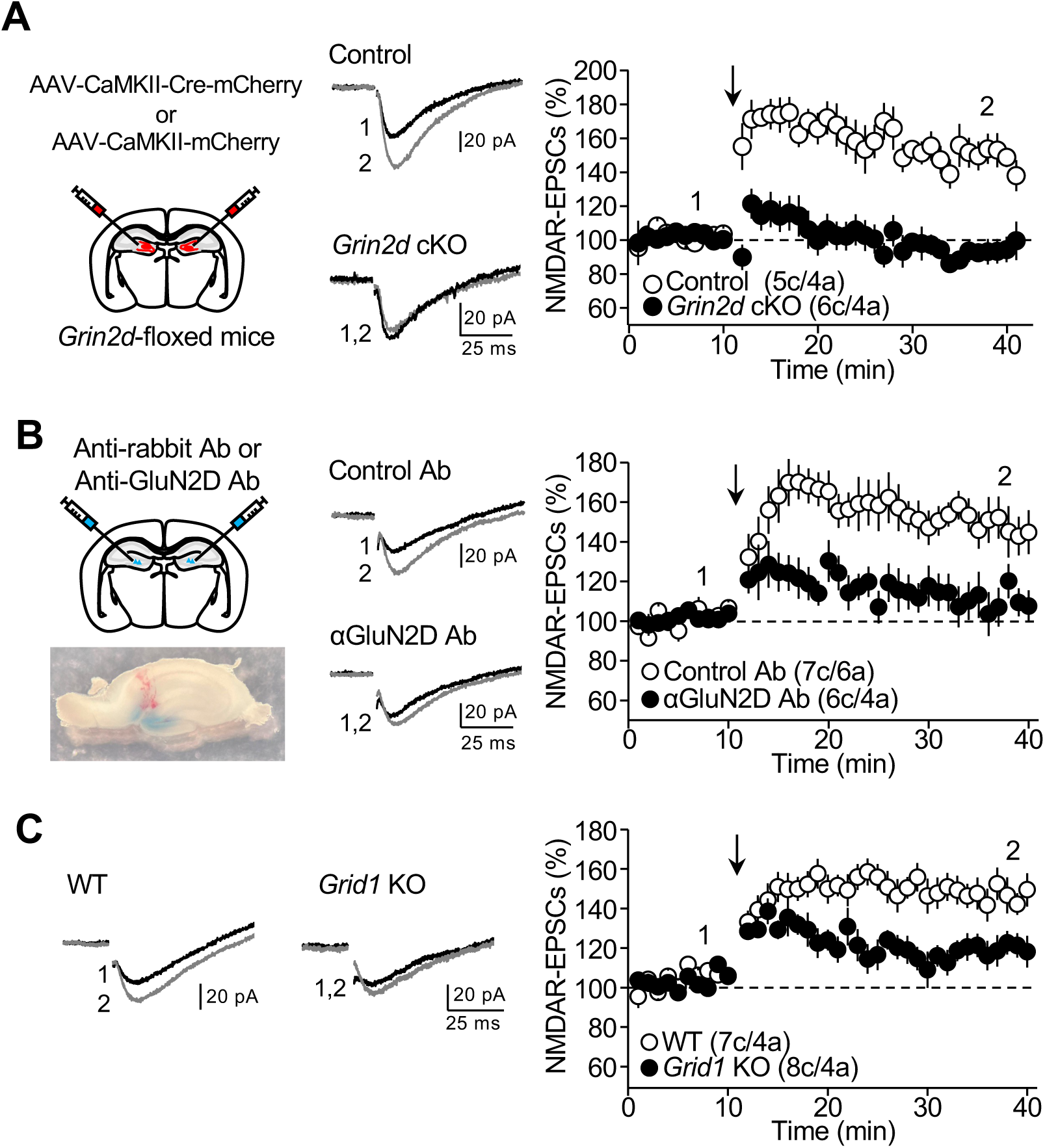
NMDAR-LTP is likely due to the recruitment of extrasynaptic GluN2D-containing NMDARs and interaction with GluD1 receptors. (A) *Grin2d^f^*^l/fl^ mice were injected with AAV_5_-CamKII-mCherry (Control) or AAV_5_-CamKII-mCherry-Cre (*Grin2d* cKO). NMDAR-LTP was abolished in *Grin2d* cKO compared with control mice (Control: 149.5 ± 6.0 %, p < 0.01, n = 5, paired t-test; cKO: 92.5 ± 5.3 %, p = 0.12201, n = 6, paired t-test; Control *vs* cKO: p < 0.001, unpaired t-test). (B) WT mice were bilaterally injected with an anti-GluN2D antibody or control Ab into the dentate gyrus. After one hour, animals were euthanized, and slices were prepared. Injection was confirmed by the presence of methylene blue. NMDAR-LTP was abolished in mice injected with the anti-GluN2D antibody (110.4 ± 8.5 %, p = 0.2952, n = 6, paired t-test) compared with control mice (149.8 ± 8.1 %, p < 0.001, n = 7, paired t-test; Control Ab *vs* anti-GluN2D Ab: p < 0.01, unpaired t-test). (C) NMDAR-LTP was impaired in *Grid1* KO mice (KO: 117.7 ± 5.3, p < 0.05%, n = 8, Wilcoxon signed-rank test) compared with controls (Control: 147.5 ± 6.7 %, p < 0.001, n = 7, paired t-test; Control *vs* KO: p < 0.05, Mann-Whitney U test). Data are presented as mean ± s.e.m.

### Extrasynaptic GluN2D-containing NMDARs are recruited at MPP-GC synapses during NMDAR-LTP by lateral diffusion

The finding that GluN2D antagonism reduced NMDAR-LTP expression without significantly affecting single-pulse-evoked synaptic transmission suggests that GluN2D-containing NMDARs are recruited to the synapse during LTP. NMDARs can undergo rapid lateral surface diffusion (*64–67*), a mechanism implicated in activity-dependent NMDAR plasticity (*27, 63, 68*). To test whether lateral diffusion mediates NMDAR-LTP at MPP-GC synapses, we artificially immobilized surface NMDARs using a cross-linking strategy (*69, 70*), in which an antibody directed against an extracellular epitope of GluN2D was injected into the dentate gyrus of wild-type mice. NMDAR-LTP was abolished in hippocampal slices (prepared one hour after injection) from mice injected with the anti-GluN2D antibody but not with a control antibody (**Figure 4B**). Importantly, the anti-GluN2D antibody infusion did not alter tonic activation of GluN2D-containing NMDARs, suggesting that NMDAR function was not significantly affected (**Figure S3**). These results support lateral diffusion of extrasynaptic GluN2D-containing NMDARs to the synapse as a key mechanism underlying NMDAR-LTP at MPP-GC synapses.

NMDARs are retained at the synapse by diverse mechanisms that involve GluN2 subunits interacting with postsynaptic scaffolding proteins in the PSD (*68, 71*). Among these proteins, PSD-95 and SAP102 dock NMDARs by binding directly to the C-terminal PDZ-binding motifs of GluN2 subunits (*72, 73*). Unlike GluN2A and GluN2B subunits, the intracellular C-terminus of GluN2D lacks PDZ-binding motifs (*1, 2*) or the sequence that allows GluN2B to interact with CaMKII⍺ (*74*), suggesting that another mechanism could retain GluN2D-containing NMDARs at the synapse following LTP induction. The non-canonical GluD1 ionotropic glutamate receptors are highly expressed in the molecular layer of the dentate gyrus (*75–77*) and interact with NMDARs (*78*). Moreover, binding of presynaptic neurexin–cerebellin complexes to postsynaptic GluD1 (*78–80*) enhances NMDAR-mediated transmission at several synapses (*81, 82*), including MPP-GC synapses (*83*). To examine whether GluD1 receptors could play a role in NMDAR-LTP at MPP-GC synapses, we tested *Grid1* KO mice. LTP was significantly reduced in these mice compared with wild-type littermates (**Figure 4C**), strongly suggesting that GluD1 is required for normal NMDAR plasticity at these synapses and could contribute to retaining GluN2D-containing receptors at the synapse.

### Deleting GluN2D from dentate gyrus excitatory neurons impaired object location memory but not object recognition memory

Lastly, we examined the behavioral consequences of genetic deletion of *Grin2d* from dentate gyrus excitatory neurons using a conditional knockout approach. To this end, we bilaterally injected a Cre-expressing virus under the CaMKII promoter (AAV_5_-CaMKII-mCherry-Cre or AAV_5_-CaMKII-mCherry as a control) into the dorsal and ventral dentate gyrus of 3–4-month-old *Grin2d^fl/fl^* (**Figure 5A**). We confirmed that these viruses targeted excitatory GCs but not inhibitory interneurons, consistent with a recent study from our group (*84*). Two weeks after injection, animals were tested in this order: open field test (OFT), object location memory (OLM), object recognition memory (ORM), and Elevated Plus Maze (EPM). During OLM training (**Figure 5B**), there was no difference in preference for either of the two objects between *Grin2d* cKO and Control mice (**Figure 5C**). During the testing phase, Control mice showed a preference for novel object location. In contrast, *Grin2d* cKO mice performed worse than Controls, as indicated by significantly lower preference scores and a lower pass rate (**Figure 5D**). No difference in total object exploration time was detected between Control and *Grin2d* cKO mice during training or testing (not shown). In a separate study, we verified that expressing Cre alone does not significantly affect OLM (*84*). For the ORM test, we used the same scoring scheme and inclusion criteria as for OLM, except that during testing, one object was replaced rather than moved (**Figure 5E**). During training, we found no difference in preference for the would-be-replaced object between *Grin2d* cKO and Control mice (**Figure 5F**). During the testing phase, there was no difference in performance between the two groups of mice as measured by either preference score or the pass rate (**Figure 5G**). Lastly, we found no significant differences between *Grin2d* cKO and Control mice in the OFT and EPM, which test locomotion and anxiety-like behaviors (**Figure S4**). Altogether, these results support that GluN2D-containing NMDARs in dentate gyrus excitatory neurons are essential for dentate gyrus-dependent forms of memory, including spatial memory, but not for object recognition memory, locomotion, or anxiety-like behaviors.

**Figure 5.**
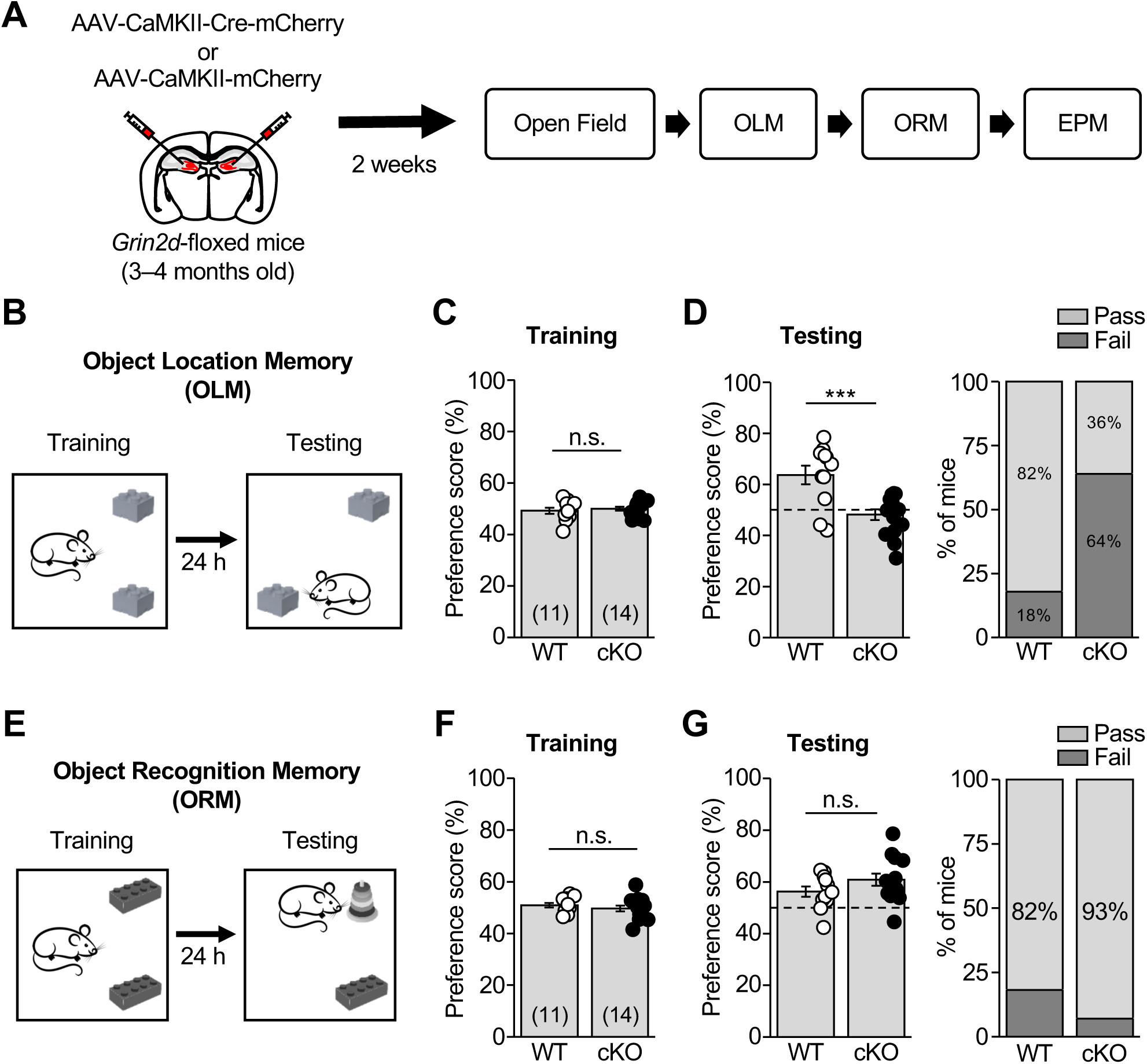
Deleting *Grin2d* from dentate gyrus excitatory neurons (*Grin2d* cKO) impaired object location memory (OLM) but not object recognition memory (ORM). **(A)** Schematic diagram illustrating the experimental timeline and strategy for conditional and selective knockout of the *Grin2d* gene from dentate gyrus excitatory neurons in adult mice (*Grin2d* cKO). All behavioral tests were performed at least 2 weeks after viral injections. **(B)** Schematic diagram illustrating OLM test. Training duration: 10 min; Test duration: 5 min; Retention interval: 24 h. **(C)** Control and *Grin2d* cKO mice did not differ significantly in preference score during training (Control: 49.2 ± 1.2 %, n = 11; cKO: 50.0 ± 0.8 %, n = 14; Control *vs* cKO: p = 0.56526, unpaired t-test). **(D)** *Grin2d* cKO mice showed impaired OLM, with a reduced preference for the moved object during testing compared with control mice (Control: 63.7 ± 3.6 %, n = 11; cKO: 48.2 ± 2.1 %, n = 14; Control *vs* cKO: p < 0.0001; unpaired t test). The dashed horizontal line indicates the threshold for passing the OLM test (mean preference score > 52 %). **(E)** Schematic diagram illustrating ORM test. Training duration: 10 min; Test duration: 5 min; Retention interval: 24 h. **(F–G)** Control and *Grin2d* cKO mice did not differ significantly in preference score during training (Control: 50.9 ± 0.91 %, n = 11; cKO: 49.7 ± 1.1 %, n = 14; Control *vs* cKO: p = 0.41488, unpaired t-test) or during testing (Control: 56.2 ± 2.0 %, n = 11; cKO: 60.9 ± 2.3 %, n = 14; Control *vs* cKO: p = 0.3117, unpaired t-test). The dashed horizontal line indicates the threshold for passing the ORM test (mean preference score > 52 %). Data are presented as mean ± s.e.m. Numbers in parentheses refer to the number of animals.

## DISCUSSION

In this study, using selective GluN2D antagonists and a conditional *Grin2d* KO approach, we report that dentate GCs express tonically active extrasynaptic GluN2D-containing NMDARs that can regulate GC firing. In addition, physiologically relevant presynaptic and postsynaptic burst activity induce LTP of NMDAR-mediated transmission at MPP-GC synapses. This potentiation is likely mediated by lateral diffusion of GluN2D-NMDARs and facilitated by GluD1 receptors (**Figure S5**). Lastly, GluN2D-containing NMDARs in excitatory neurons, including GCs, are essential for normal dentate gyrus-dependent memory. Altogether, these findings indicate that GluN2D-containing NMDARs play important roles in hippocampal function, most likely by regulating GC firing and mediating synaptic plasticity.

The GluN2D subunit is highly expressed during embryonic and early postnatal development, and its forebrain expression declines sharply with maturation (*4, 5, 11*). In the mature brain, GluN2D-containing NMDA receptors are more characteristic of GABAergic interneurons than of glutamatergic principal neurons. Pharmacological studies show that GluN2D-containing NMDARs in inhibitory interneurons are primarily extrasynaptic and mediate tonic currents that facilitate interneuron activity (*18, 24, 25, 48, 50, 51*). There is little evidence that these receptors play a role in the excitatory neurons of the mature cortex and hippocampus. Our in situ hybridization results revealed *Grin2D* mRNA expression in GCs, consistent with a previous human tissue study (*26*). Moreover, we found that excitatory GCs express functional GluN2D-containing NMDARs that are tonically active in brain slices from adult rats and mice, as indicated by reductions in both holding current and induced-firing activity mediated by the specific GluN2D antagonists DQP-1105 and QNZ46. Remarkably, this effect was abolished in *Grin2d* cKO mice, supporting the key role of GluN2D-containing NMDARs in regulating GC firing. Ambient glutamate likely activates these receptors, as has been reported for the tonic activation of NMDARs in other neurons (*44–48*).

GluN2D-containing NMDARs also mediate basal synaptic transmission in several brain areas, including hippocampal inhibitory interneurons (*14, 17, 18*), subthalamic neurons (*19*), dopaminergic neurons in the substantia nigra (*23*), and inferior colliculus (*20*). In dentate GCs, however, we found that these receptors do not contribute significantly to basal transmission but can be activated by brief repetitive stimulation of MPP inputs and recruited to the synapse after LTP induction. Previous studies used high-frequency, bulk stimulation of perforant path axons to elicit NMDAR-LTP in GCs of rat hippocampal slices (*27, 41*). We found that a more physiological BTDP protocol, which induces LTP of NMDAR-mediated synaptic transmission at other synapses (*56–58*), also triggered NMDAR LTP at MPP-GC synapses in both mice and rats. This induction protocol mimics *in vivo* activity in the entorhinal cortex (*59, 60*) and the dentate gyrus (*61, 62*). Our findings are consistent with a previous study (*27*) that used early antagonists, such as PPDA and UBP141, which exhibit modest selectivity for GluN2D-containing NMDARs (*85, 86*) and whose effects were not validated with alternative approaches. Since then, highly selective GluN2D antagonists have been developed. In our study, we used two of these antagonists, as well as *Grin2d* cKO mice, to demonstrate that NMDAR-LTP at MPP-GC synapses is mediated by the recruitment of GluN2D-containing receptors.

An increase in synaptic NMDARs during LTP can occur via exocytosis or lateral diffusion (*63*). Our results using a cross-linking approach support lateral diffusion of extrasynaptic receptors, in line with previous studies showing rapid lateral diffusion of NMDARs at the synapse (*65, 66, 69, 70*); for a recent review, see (*68*). NMDARs are retained at the synapse primarily through scaffolding proteins (MAGUKs) that bind the C-terminal PDZ binding motif of GluN2 subunits (*68, 71–73*), which GluN2D lacks (*1, 2*). Thus, anchoring of GluN2D-containing receptors at the synapse could involve interactions with transmembrane proteins, such as ion channels or neurotransmitter receptors, as well as transsynaptic complexes (*68*). We hypothesize that GluD1 receptors, which are highly expressed in the molecular layer of the dentate gyrus (*75–77*), interact with NMDARs (*78*), and form transsynaptic complexes with presynaptic neurexin and cerebellin (*78–80*), thereby contributing to the retention of GluND2-containing NMDARs at the MPP-GC synapse following LTP induction. In support of this hypothesis, we found that NMDAR-LTP was significantly reduced in *Grid1* KO mice. Previous studies have shown that binding of presynaptic neurexin–cerebellin complexes to postsynaptic GluD1 enhances NMDAR-mediated transmission at several synapses, including MPP-GC synapses (*81–83*). Cerebellin (Cbl) 1 and 4 mRNAs are highly expressed in the entorhinal cortex neurons that project to the molecular layer of the dentate gyrus via perforant path axons (*87, 88*). Moreover, Cbl4 can be secreted from these axons during repetitive activity to form a transsynaptic complex with presynaptic neurexin and postsynaptic neogenin-1, and, remarkably, this complex is essential for NMDAR-dependent AMPAR-LTP at MPP-GC synapses (*89*). It is tempting to postulate that a similar mechanism, though likely involving a neurexin/Cbln4/GluD1 transsynaptic complex, is required for NMDAR-LTP. In the hippocampus, GluD1 colocalizes with and functionally interacts with the type 5 metabotropic glutamate receptor (mGluR5) (*90*). Thus, GluD1 could potentiate NMDAR-mediated transmission through multiple mechanisms.

A similar protocol we used to trigger NMDAR-LTP also induces LTP of AMPAR-mediated transmission at entorhinal-GC synapses (*91, 92*). Whether long-term NMDAR plasticity can be induced without inducing AMPAR-LTP remains to be tested. If so, NMDAR-LTP at MPP-GC synapses could mediate metaplasticity (*63*) –i.e., changes in the inducibility of AMPAR-LTP or LTD that rely on NMDAR activation. Intriguingly, reversing the burst order (post-pre), which induces NMDAR-LTD at other synapses in the midbrain (*56*) and in the CA3 area (*57*), had no long-lasting effect at MPP-GC synapses. This observation is consistent with the low contribution of GluN2D-containing NMDARs to basal transmission at these synapses and supports the notion that synaptic learning rules and NMDAR plasticity mechanisms differ across synapse types (*58, 63, 93*). Whether depotentiation of NMDAR-mediated transmission at MPP-GC synapses occurs spontaneously or can be modulated by activity or neuromodulators remains to be determined.

Because GCs provide a major synaptic input to CA3 pyramidal neurons, GC spike generation is likely to play a significant role in hippocampal information processing (*94*). Our results show that GluN2D-containing NMDARs can facilitate GC firing through tonic activation of extrasynaptic receptors and by mediating NMDAR-LTP at MPP-GC synapses. Both processes, which are blocked in *Grin2d* cKO, could contribute to the memory deficit observed in these mice. LTP induction enhanced GC firing in response to brief bursts of MPP synaptic input. Although NMDAR-mediated currents are small at the resting membrane potential, current flow through the channel can be amplified by depolarization during burst-like excitatory activity, as occurs *in vivo* (*63*). The slow decay of NMDAR-EPSPs enables temporal summation, which can sustain excitation and drive neuronal firing during repetitive synaptic activity, as recently reported for GluN2D-containing NMDARs in the inferior colliculus (*20*). Thus, potentiation of NMDAR-mediated transmission at MPP-GC synapses may significantly contribute to the integrative properties of GCs and their recruitment into neuronal ensembles. Lastly, dysregulated tonic activation of GluN2D-containing NMDARs in GCs could contribute to brain disorders, including epileptic discharge (*95*) and neuronal loss (*8*).

## MATERIALS AND METHODS

### Experimental Model and Subject Details

Adult Sprague-Dawley rats (Charles River), C57BL/6J (Jackson Labs), *Grin2d* KO, floxed-*Grin2d* (*Grin2d*^fl/fl^) mice, and *Grid1* KO mice of either sex were used for electrophysiological and behavioral experiments. The *Grin2d* KO mice were generated using *Grin2d*^tm1a(EUCOMM)Wtsi^ (KO-first allele, GluN2D KO) mice (Wellcome Trust Sanger Institute). The *Grin2d*^fl/fl^ mice were generated by removing the reporter cassette by crossing *Grin2d*^tm1a(EUCOMM)Wtsi^ mice with B6-SJL-Tg(ACTFLPe)9205Dym/J line. *Grid1* KO mice(*96*) were a gift from Dr. Jian Zuo (St. Jude Children’s Research Hospital). All animals were group-housed on a standard 12-hr light/12-hr dark cycle and had free access to food and water. Animal handling and use were conducted in accordance with protocols approved by the Animal Care and Use Committees of Albert Einstein College of Medicine, Texas A&M University, and Virginia Polytechnic Institute and State University, as well as with the National Institutes of Health guidelines.

### Stereotaxic injection

C57BL/6J and *Grin2d*^fl/fl^ mice were anesthetized with isoflurane (up to 5% for induction and 1%–3% for maintenance) and placed on a stereotaxic frame. AAV_5_-CamKII-mCherry (Control) or AAV_5_-CamKII-Cre-mCherry (University of Pennsylvania Vector Core) was injected (1 μL at 0.1 μL/min) bilaterally into the dentate gyrus (2.06 mm posterior to bregma, ±1.5 mm lateral to bregma, 1.65 mm ventral from dura) of 4–5-week-old *Grin2d*^fl/fl^ mice. Slices for electrophysiological recording were prepared 10–21 days after surgery. For GluN2D cross-linking experiments in C57BL/6J, the control group received 1 μL of anti-rabbit Alexa 568 (control IgG, 1/5, Invitrogen A11008), while the GluN2D-cross-link group received 1 µg of polyclonal rabbit anti-GluN2D subunit (Alomone Labs, cat #AGC-020), both diluted in PBS with 1% methylene blue (1 µL final volume). Slices for electrophysiology were prepared 1 hour after surgery, and injection was confirmed by the presence of methylene blue.

### Hippocampal slice preparation

Acute transverse hippocampal slices were prepared from Sprague-Dawley rats, C57BL/6J (P21-P35), *Grin2d*^fl/fl^, *Grin2d* KO, and *Grid1* KO mice (P28–P42); 300- or 400-μm thick for mice and rats, respectively. Briefly, rat hippocampi were isolated and cut with a VT1200S microslicer (Leica Microsystems Co.) in a solution containing (in mM): 215 sucrose, 2.5 KCl, 26 NaHCO_3_, 1.6 NaH_2_PO_4_, 1 CaCl_2_, 4 MgCl_2_, 4 MgSO_4_, and 20 D-glucose. For mice, hippocampi were dissected and sliced in an ice-cold cutting solution containing (in mM): 110 choline, 2.5 KCl, 25 NaHCO_3_, 1.25 NaH_2_PO_4_, 0.5 CaCl_2_, 7 MgCl_2_, 25 D-glucose, 11.6 sodium L-ascorbate, and 3.1 sodium pyruvate. Slices were then transferred to a chamber at room temperature with extracellular artificial cerebrospinal fluid (ACSF) containing (in mM): 124 NaCl, 2.5 KCl, 26 NaHCO_3_, 1 NaH_2_PO_4_, 2.5 CaCl_2_, 1.3 MgSO_4_, and 10 D-glucose. Slices were kept in this chamber for at least 30 min before recording. All solutions were equilibrated with 95% O_2_ and 5% CO_2_ (pH 7.4).

### Electrophysiology

All experiments, unless otherwise stated, were performed at 28 ± 1°C in a submersion-type recording chamber perfused at ∼2 mL min^−1^ with ACSF supplemented with 10 μM NBQX and 100 μM picrotoxin to block AMPA/Kainate and GABA_A_ receptors, respectively. Whole-cell patch-clamp recordings were performed using a Multiclamp 700A amplifier (Molecular Devices). GCs were voltage clamped at – 45 mV (unless otherwise stated) using patch-type pipette electrodes (∼3–4 MΩ) containing (in mM): 135 K-Gluconate, 5 KCl, 0.1 EGTA, 0.04 CaCl_2_, 5 NaOH, 5 NaCl, 10 HEPES, 5 MgATP, 0.4 Na_3_GTP, and 10 D-glucose, pH 7.2 (280−290 mOsm). Voltage-clamp recordings of GCs (Vh = +40 mV) shown in Figure 2A–B, 2E were performed using a Cs-based internal solution containing (in mM): 131 cesium gluconate, 8 NaCl, 1 CaCl2, 10 EGTA, 10 D-glucose, and 10 HEPES, pH 7.2 (285–290 mOsm). Series resistance (∼7–30 MΩ) was monitored throughout all experiments using a −5 mV, 80 ms voltage step, and cells exhibiting a significant change in series resistance (>20%) were excluded from analysis. Action potentials (APs) were elicited by injecting 500-ms depolarizing current steps in the presence of 10 μM NBQX and 100 μM picrotoxin to block AMPAR- and GABA_A_-mediated synaptic transmission, respectively. To stimulate MPP inputs to GCs, a stimulating patch-type pipette was placed in the middle third of the medial molecular layer of the dentate gyrus. To elicit synaptic responses, monopolar square-wave voltage or current pulses (100–200 μs pulse width, 4–25 V or 20–100 μA) were delivered through a stimulus isolator (Isoflex, AMPI, or Digitimer DS2A-MKII) connected to a broken-tip (∼10–20 μm) stimulating patch-type micropipette filled with ACSF. Typically, stimulation was adjusted to obtain comparable-magnitude synaptic responses across experiments, e.g., 40–100 pA NMDAR-EPSCs (V_h_ = −45 mV). Burst-timing NMDAR plasticity was typically induced in current-clamp mode (V_rest_ ∼ –60 to – 70 mV) by pairing presynaptic bursts (6 pulses at 50 Hz) with postsynaptic bursts of action potentials (5 action potentials at 100 Hz) delivered at 10-ms intervals, repeated 100 times at 2 Hz. To examine synaptically-driven activation of GCs (Figure 3G–H), we performed whole-cell current-clamp recordings from these cells (Vm –60 mV) and measured the probability of AP firing in response to MPP stimulation consisting of 5 pulses delivered at 50 Hz, across different stimulus intensities (10 V, 20 V, 30V). We used 3 sweeps for each stimulation intensity and calculated the average number of AP/train. We monitored MPP-evoked firing before LTP induction and at 15–20 min after LTP induction. To isolate NMDAR-EPSPs, experiments were performed in the continuous presence of 10 µM NBQX, 100 µM picrotoxin, and 20 µM SCH50911 (GABA_B_ receptor antagonist). NMDA-evoked responses (Figure 2E) were triggered by puffing 1 mM NMDA and 100 µM glycine (20 ms, 5–6 PSI) using a Picospritzer III (Parker) connected to a patch pipette (3 MΩ). The tip of the puffer pipette was in the MML < 30 μm deep from the surface of the slice (∼200 µm from the granule cell soma). Recordings were obtained at +40 mV in the presence of 1 μM Tetrodotoxin, 10 μM NBQX, and 100 μM picrotoxin. Electrophysiological data were acquired at 5 kHz, filtered at 2.4 kHz, and analyzed using custom software for IgorPro 7.01 (Wavemetrics Inc.). The magnitude of LTP was determined by comparing 10-min baseline responses with responses 20–30 min after LTP induction. Averaged traces include 20 consecutive individual responses. All experiments were performed in an interleaved fashion – i.e., control experiments were performed every test experiment on the same day.

### Behavior analysis

Mice were extensively handled before behavioral analysis. Testing was conducted in a Plexiglas box (width: 42 cm, length: 42 cm, height: 42 cm) placed in a dimly lit room with clearly visible contextual cues (black-on-white patterns). Mice were tested in the open field for 10 min on day 1. On days 2 and 3, mice performed the object location memory test, and on days 4 and 5, the object recognition memory test. On day 6, mice were tested in the elevated plus-maze. For the object location and object recognition memory tests, mice underwent a 10-min training session, a 24-h retention interval during which they were returned to the home cage, and a 5-min test session. The objects were plastic toys (approximately 5 cm in width, 5 cm in length, 7 cm in height) and were cleaned with 70% ethanol between sessions. Experiments were video-recorded, and object exploration times (nose in contact with the object or sniffing the object at < 1 cm) were measured by a blinded observer. Experimenters were blinded to the condition during both performance and analysis. The preference score (%) was calculated as [(exploration of novel object/total object exploration)*100]. Mice with a preference score > 52 % passed the test. Data were analyzed using OriginPro software (OriginLab).

#### Open Field Test (OFT)

Mice were placed individually in the center of the arena and allowed to explore freely for 10 min. Using the automated video-tracking software EthoVision (Noldus), the total distance traveled and the time spent in the arena center (18 cm x 18 cm) were measured during the first 5 min.

#### Object Location Memory (OLM)

During the training phase, animals were allowed to explore two objects for 10 min and were then returned to their home cage for 24 h. After this delay, the animals were returned to the arena containing the objects used in the training phase. One object was placed in the location it occupied during the training phase, while the other was placed in a new location within the arena. Object exploration was recorded for 5 min, after which the animals were removed and returned to their home cage.

#### Object Recognition Memory (ORM)

During the training phase, animals were allowed to explore two distinct objects for 10 min and were then returned to their home cage for 24 h, during which the arena and objects were cleaned. Animals were then returned to the arena with an identical copy of one familiar object and a novel object, and were allowed to explore freely for 5 min. At the end of this recognition phase, animals were removed and returned to their home cage.

#### Elevated-Plus Maze (EPM)

Mice were placed individually in the center area, facing an open arm, and allowed to explore the apparatus freely for 5 min. Using the automated video-tracking software EthoVision (Noldus), the total distance traveled, and the time spent in open or closed arms were measured.

### Post-hoc analysis of AAV expression

At the end of the EPM, viral expression was verified *post hoc,* and mice showing less than 75% viral expression were excluded. Mice were anesthetized with isoflurane (3–5%) and transcardially perfused with 4% paraformaldehyde (PFA) in 0.1 M phosphate-buffered saline (PBS). 50 µm-thick coronal brain sections were prepared with a Leica Microslicer (VT1000S), stained with DAPI (1:1000) to label cell nuclei, and mounted with Prolong Diamond antifade mounting reagent (ThermoFisher) onto microscope slides. All images in this study were acquired on a Zeiss LSM 880 Airyscan confocal microscope with Super-Resolution and ZEN (black edition) software, using a 25X oil-immersion objective.

### Fluorescence in situ hybridization

Mice were euthanized by overdose with 10% isoflurane using the open-drop method. The mice were decapitated, and brains were dissected, embedded in Optimal Cutting Temperature (OCT) compound, and rapidly frozen by submersion in a 2-methylbutane bath on dry ice. Twenty-micron cryosections were mounted on slides and then processed for single-molecule fluorescence in situ hybridization (FISH) according to the manufacturer’s instructions in the RNAscope HiPlex kit (Advanced Cell Diagnostics). Probes used included: *Grin2d* (425951-T5)*, Grin1 (*431611-T7), and *parvalbumin* (421931-T8), as well as the manufacturer’s negative and positive controls. Z-stack images (15 slices, 0.3 µm interval) were acquired using a Leica Stellaris SP8 confocal microscope with a 63X (1.4 NA) objective. RNAscope probe fluorophores were excited using the 488 nm, 647 nm, and 750 nm laser lines of the white light laser, and DAPI was imaged using a 405 nm laser. Laser power and gain settings were such that the negative control had ≤ 2 particles visible per cell. Images were exported as 16-bit TIFF files, and maximum-intensity projections were generated in Fiji.

### Reagents

Reagents were bath-applied after dilution of stock solutions in ACSF, stored at 4°C or –20°C, prepared in water or DMSO, depending on the manufacturer’s recommendations. The final DMSO concentration was < 0.01% of the total volume. DQP-1105, QNZ46, D-APV, NMDA, SCH50911, and glycine were purchased from Tocris Biosciences. NBQX and Tetrodotoxin were purchased from Cayman Chemical Co., and picrotoxin was purchased from Hello Bio. Methylene blue, and all salts used to prepare ACSF and internal solutions were purchased from Sigma-Aldrich.

### Quantification and Statistical Analysis

Statistical analysis was performed using OriginPro software (OriginLab). Normality of distributions was assessed using the Shapiro-Wilk test. For normal distributions, unpaired and paired two-tailed Student’s t tests were used to assess between-group and within-group differences, respectively. Parameters across more than two groups were compared using a one-way ANOVA followed by Tukey’s post hoc analysis or two-way ANOVA followed by Tukey’s post hoc test when more than one parameter was included in the comparison. For non-normal distributions, the nonparametric Wilcoxon signed-rank test and Mann-Whitney U test were used. Statistical significance was set at p < 0.05 (*** indicates p < 0.001, ** indicates p < 0.01, and * indicates p < 0.05). All values are reported as the mean ± s.e.m.

## Acknowledgments

We thank Drs. Steve Traynelis (Emory School of Medicine), Pierre Paoletti (École Normale Supérieure de Paris), and all the Castillo lab members for their constructive feedback. We also thank Dr. Laurent Groc (University of Bordeaux) for helpful advice regarding the cross-linking experiments.

## Funding

This work was supported by NIH grants R01 MH116673, R01 MH125772, and R01 NS113600 to P.E.C.; R01 MH116003, R01 NS118731, and R01 NS133338 to S.M.D.; and R01 NS105804 to S.A.S. C.B. was partially supported by a Junior Investigator Neuroscience Research Award (Einstein), an American Epilepsy Society Postdoctoral Research Fellowship, and a RFK-IDDRC Pilot Grant (NIH P50 HD105352). A.R.R. was partially supported by the Brain & Behavior Research Foundation Young Investigator Award and the Ford Foundation Postdoctoral Fellowship. K.N. was partially supported by Postdoctoral Research Fellowships from the American Epilepsy Society, the Fondation pour la Recherche Médicale, and the Fondation Bettencourt Schueller. M.C. was supported by NIH R25GM104547. We thank the Wellcome Trust Sanger Institute Mouse Genetics Project (Sanger MGP) and its funders for providing the mutant mouse line (*Grin2d*^tm1a(EUCOMM)Wtsi^). Funding and associated primary phenotypic information may be found at www.sanger.ac.uk/mouseportal. We thank Dr. Jian Zuo (St. Jude Children’s Research Hospital) for sharing *Grid1* KO mice. Confocal images were obtained at the Einstein Imaging Corle (supported by The Rose F Kennedy Intellectual Disabilities Research Center U54 HD090260 and shared instrument grant NIH 1S10OD25295 to Konstantin Dobrenis).

## Author contributions

C.B., A.R.R., K.N., S.A.W., and P.E.C. designed the experiments. C.B., A.R.R., L.B., K.N., and M.C. performed and analyzed electrophysiological recordings. S.A.W. performed RNAscope experiments. K.N. and G.P.S. performed puff experiments under the supervision of S.M.D. C.B. performed stereotaxic injections and behavior. C.B. and P.E.C. wrote the manuscript, and all authors read and edited the manuscript.

## Competing interests

The authors declare no competing interests.

## Notes

### Competing Interest Statement

The authors have declared no competing interest.

### Summary of Updates

The manuscript was updated with new results.

