## Supplemental Figures for "GluN2D-containing NMDA receptors support dentate granule cell excitability, synaptic plasticity, and memory"

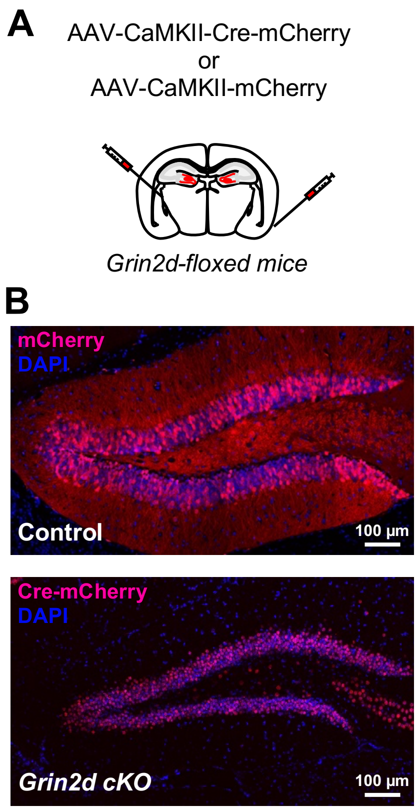


**Fig S1. *Grin2d* conditional knockout.**

**(A)** *Grin2d*$^{fl/fl}$ mice were injected with AAV-CamKII-mCherry (Control) or AAV-CamKII-mCherry-Cre (*Grin2d* cKO).

**(B)** Confocal images showing expression of mCherry and Cre-mCherry in the dentate gyrus of *Grin2d*$^{fl/fl}$ mice.

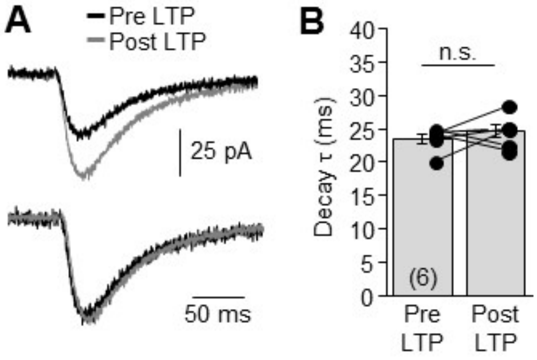


**Fig S2. NMDAR-EPSC decay was not changed after NMDAR-LTP induction.**

**(A)** Top, representative NMDAR-EPSCs before and after LTP induction. Bottom, peak-scaled traces.

**(B)** EPSC decay was unchanged after LTP induction (Pre LTP: 23.5 ± 0.7 ms; Post LTP: 24.6 ± 1.0 ms; $p = 0.4323$, $n = 6$, paired t test).

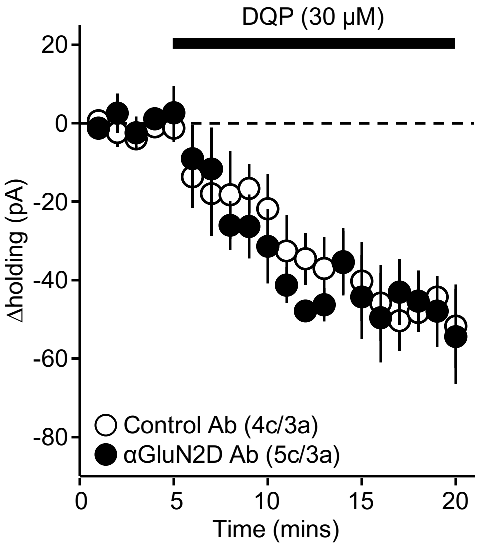


**Fig S3. Tonic activity of GluN2D-containing NMDARs was not affected by anti-GluN2D antibody infusion into the dentate gyrus.**

Bath application of the GluN2D antagonist DQP-1105 (30 μM for 15 min) similarly decreased the GC holding current in mice injected with anti-GluN2D antibody (-47.7 ± 8.9 pA, $p < 0.01$, $n = 5$, paired t-test) and control antibody (-43.7 ± 5.8 pA, $p < 0.01$, $n = 4$, paired t-test). Voltage-clamp recordings of GCs were performed at +40 mV, in the presence of 10 μM NBQX and 100 μM picrotoxin.

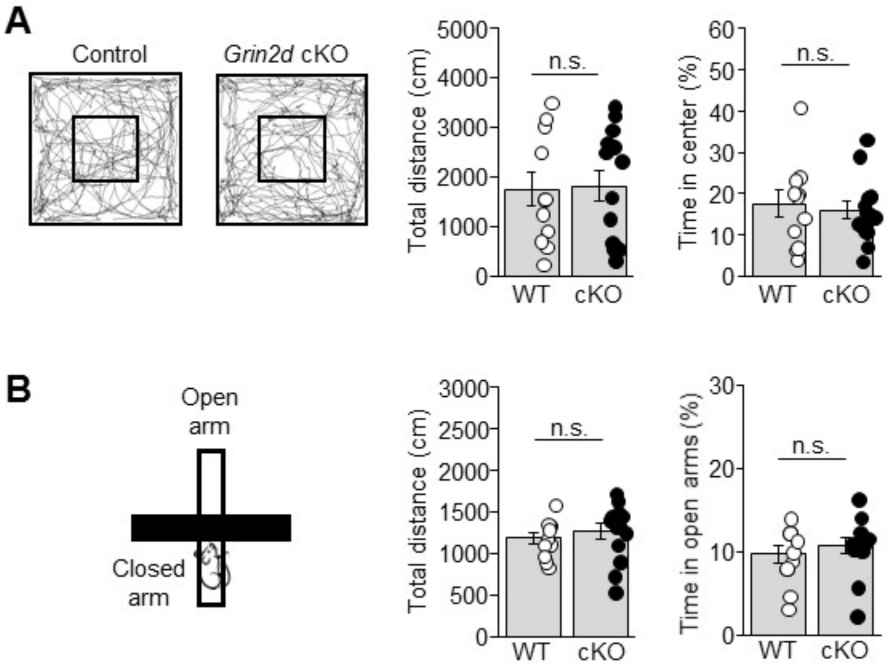


**Fig S4. Deleting *Grin2d* from dentate gyrus excitatory neurons (*Grin2d* cKO) did not affect locomotion and anxiety-like behaviors.**

**(A)** *Left,* Representative track maps of a Control and a GC *Grin2d* cKO mouse in the open field. *Right,* Control and *Grin2d* cKO mice did not differ significantly in total distance travelled (Control: 1742 ± 345 cm, n = 11; cKO: 1811 ± 305 cm, n = 14; Control *vs* cKO: p = 0.88221, unpaired t-test) or time spent in center of the arena (Control: 18 ± 3.2 %, n = 11; cKO: 16 ± 2.1 %, n = 14; Control *vs* cKO: p = 0.68331, unpaired t-test).

**(B)** *Left,* Schematic diagram illustrating the elevated-plus maze (EPM) test. *Right,* Control and *Grin2d* cKO mice did not differ significantly in total distance travelled (Control: 1192 ± 68 cm, n = 11; cKO: 1267 ± 97 cm, n = 13; Control *vs* cKO: p = 0.54711, unpaired t-test) or time spent in the open arms (Control: 10 ± 1.0 %, n = 11; cKO: 11 ± 1.0 %, n = 13; Control *vs* cKO: p = 0.48691, Mann-Whitney U test).

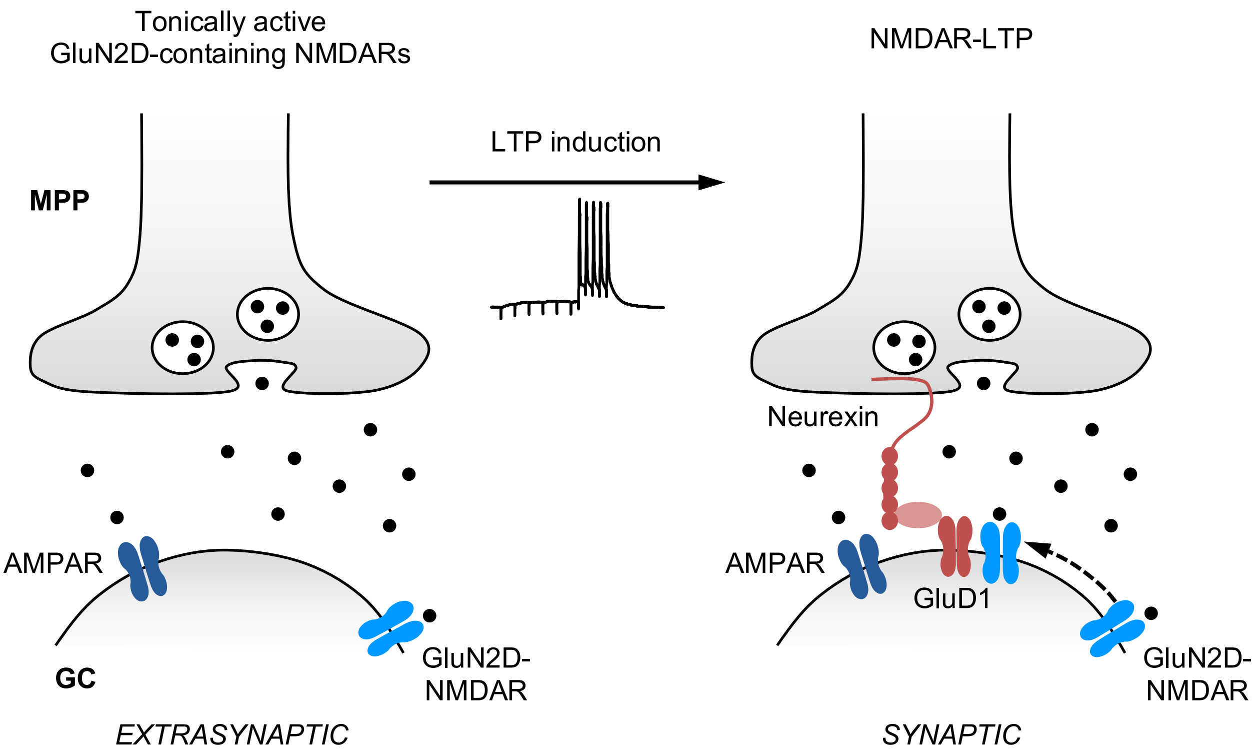


**Fig S5. Role of GluN2D-containing NMDA receptors in the dentate gyrus**

Tonically active extrasynaptic GluN2D-containing NMDA receptors (GluN2D-NMDARs) regulate granule cell (GC) action potential firing. After induction of burst timing-dependent plasticity (BTDP) of NMDAR-mediated transmission at medial perforant path (MPP)-GC synapses, GluN2D-NMDARs localize at the synapse via lateral diffusion. GluD1 receptors contribute to the retention of GluN2D-containing NMDARs at the MPP-GC synapse following LTP induction, likely by forming transsynaptic complexes with presynaptic neurexin and cerebellin.
